# HCMV glycoprotein B subunit vaccine efficacy was mediated by non-neutralizing antibody effector functions

**DOI:** 10.1101/246884

**Authors:** Cody S. Nelson, Tori Huffman, Eduardo Cisneros de la Rosa, Guanhua Xie, Nathan Vandergrift, Robert F. Pass, Justin Pollara, Sallie R. Permar

## Abstract

Human cytomegalovirus (HCMV) is the most common congenital infection worldwide, frequently causing hearing loss and brain damage in afflicted infants. A vaccine to prevent maternal acquisition of HCMV during pregnancy is necessary to reduce the incidence of infant disease. The glycoprotein B (gB) + MF59 adjuvant subunit vaccine platform is the most successful HCMV vaccine tested to-date, demonstrating approximately 50% efficacy in preventing HCMV acquisition in phase II trials. However, the mechanism of vaccine protection remains unknown. Plasma from 33 gB/MF59 vaccinees at peak immunogenicity was tested for gB epitope specificity as well as neutralizing and non-neutralizing anti-HCMV effector functions, and compared to an HCMV-seropositive cohort. gB/MF59 vaccination elicited IgG responses with gB-binding magnitude and avidity comparable to natural infection. Additionally, IgG subclass distribution was similar with predominant IgG1 and IgG3 responses induced by gB vaccination and HCMV infection. However, vaccine-elicited antibodies exhibited limited neutralization of the autologous virus, negligible neutralization of multiple heterologous strains, and limited binding responses against gB structural motifs targeted by neutralizing antibodies including AD-1, AD-2, and Domain I. Interestingly, vaccinees had high-magnitude IgG responses against AD-3 linear epitopes, demonstrating immunodominance against this non-neutralizing, cytosolic region. Finally, vaccine-elicited IgG robustly bound trimeric, membrane-associated gB on the surface of transfected or HCMV-infected cells and mediated virion phagocytosis, though were poor mediators of NK cell activation. Altogether, these data suggest that non-neutralizing antibody functions, including virion phagocytosis, likely played a role in the observed 50% vaccine-mediated protection against HCMV acquisition.

**Significance:** The CDC estimates that every hour, a child is born in the United States with permanent neurologic disability resulting from human cytomegalovirus (HCMV) infection – more than is caused by Down syndrome, fetal alcohol syndrome, and neural tube defects combined. A maternal vaccine to block transmission of HCMV to the developing fetus is a necessary intervention to prevent these adverse outcomes. The gB/MF59 vaccine is the most successful tested clinically to-date, achieving 50% reduction in HCMV acquisition. This manuscript establishes the function and epitope specificity of the humoral response stimulated by this vaccine that may explain the partial vaccine efficacy. Understanding the mechanism of gB/MF59-elicited protective immune responses will guide rational design and evaluation of the next generation of HCMV vaccines.

## Introduction

Human cytomegalovirus (HCMV) is the most common congenital viral infection (1), impacting 1 out of every 150 live born infants worldwide (2). In the United States alone, this equates to 40,000 children infected annually, of whom 8,000 develop long-term disabilities including microcephaly, intrauterine growth restriction, hearing/vision loss, or neurodevelopmental delay (3, 4) – more congenital disease than all 29 newborn conditions currently screened for in the US combined (5). It is clear that preexisting maternal immunity impacts the incidence of congenital infection, as 30-40% of HCMV-seronegative women that acquire the virus during pregnancy transmit the infection to the fetus *in utero* in contrast to 1-2% following superinfection of HCMV-seroimmune women (3). Furthermore, infants born to HCMV-seropositive women are also less likely to exhibit symptoms of congenital infection at birth and can have milder neurologic sequelae (6, 7). Therefore, many hypothesize that a maternal vaccine that prevents maternal HCMV acquisition, protects against viral transmission to the infant, or reduces the severity of congenital infection is an achievable goal (8-10). Given the global burden of disease of congenital HCMV, such a vaccine remains a Tier 1 priority of the National Academy of Medicine (11).

A variety of candidate HCMV vaccine approaches have been attempted, including live attenuated virus (12, 13), glycoprotein subunit formulations (14, 15), viral vectors (16, 17), and single/bivalent DNA plasmids (18, 19). Importantly, the HCMV glycoprotein B (gB) subunit vaccine administered with MF59 squalene adjuvant demonstrated moderate (~50%) efficacy in preventing primary HCMV infection in cohorts of both postpartum (14) and adolescent women (15). Furthermore, this vaccine demonstrated a protective benefit against HCMV viremia and reduced clinical need for antiviral treatment in transplant recipients (20). gB is the viral fusogen which facilitates the fusion of viral envelope and host membrane during viral entry (21). This glycoprotein is essential for entry into all cell types, including trophoblast progenitor cells (22), and is a known target of neutralizing antibodies (23, 24). However, previous investigations have reported that gB/MF59-elicited antibodies were poorly neutralizing (25-27), raising questions about the mechanism underlying the partial gB vaccine efficacy observed in multiple clinical trials. An understanding of the gB/MF59-mediated protection is needed to rationally-design immunogens that will improve upon the partial vaccine efficacy that was achieved clinically.

Glycoprotein B is a 907-amino acid, homotrimeric glycoprotein consisting of 4 distinct structural regions: an ectodomain, a membrane-proximal region (MPER), a transmembrane domain, and a cytoplasmic domain (Fig. S1) (28, 29). Additionally, there are 5 distinct antigenic sites known to be targeted by gB-specific antibodies, identified as antigenic domain (AD) 1-5 (Fig. S1) (23, 28, 29). Antibodies against AD-1, an uninterrupted ~80 amino acid epitope, are present in virtually all infected individuals (23) and can be either neutralizing or non-neutralizing (30, 31). In contrast, AD-2 specific antibodies are present in only subset of seropositive people (23, 32), and this region consists of 2 unique linear epitopes: site 1 is perfectly conserved in all viral strains and a target of potently-neutralizing antibodies, whereas site 2 is highly variable and targeted by only non-neutralizing antibodies (32). AD-3 is located within the cytosolic domain and is a known target of exclusively non-neutralizing antibodies (33-35). Finally, AD-4 (Domain AI) and AD-5 (Domain I) are conformational, globular protein domains that were recently identified and characterized as the targets of neutralizing antibodies (23, 36-38). The antigen used for the gB/MF59 vaccine clinical trials (14, 15, 20) consisted of the full protein with two modifications to facilitate manufacture: (1) deletion of the transmembrane domain (75 amino acids) and (2) mutation of the furin protease cleavage site (39).

To identify possible mechanisms accounting for the partial protection against HCMV infection following gB/MF59 vaccination, we undertook an in-depth investigation into the characteristics and functionality of the antibody responses elicited by this gB subunit vaccine. As other groups have reported, we observed that gB/MF59 vaccine-elicited antibodies mediated limited neutralization. Here we report on our observations that gB/MF59 vaccination results in an antibody profile quite distinct from that observed in the setting of natural HCMV infection.

Furthermore, we have observed that the poorly-neutralizing, gB/MF59-elicited antibodies can mediate robust non-neutralizing antiviral responses, such as phagocytosis, leading us to propose that non-neutralizing antibody functions played a role in the observed 50% vaccine efficacy against HCMV acquisition.

## Results

### Immunogen and HCMV virion binding IgG responses/avidity

We obtained plasma samples from 33 gB/MF59 vaccinee participants in the phase 2 clinical trial conducted in a population of seronegative postpartum women (14). All subsequent studies used samples collected at peak immunogenicity (6.5 months), or the next available (not exceeding 12 months). The gB/MF59 vaccine platform has been previously reported to elicit robust titers of gB immunogen-specific IgG (39, 40). We observed similar results for this subset of vaccinees (gB/MF59), with high-magnitude plasma gB IgG binding exceeding that elicited in chronically-infected, seropositive (SP) individuals (Fig. 1A) (log_10_AUC: gB/MF59=6.32, SP=5.64; p=0.03, Wilcoxon rank sum test). In contrast, much lower IgG binding of vaccinee sera was observed against whole HCMV virions compared to seropositive individuals (Fig. 1B) (log_10_AUC: gB/MF59=0.48, SP=2.91; p<0.001, Saittertherwaite t test), likely due to naturally HCMV-elicited antibodies targeting a variety of other CMV glycoproteins and/or multiple episodes of reactivation or reinfection boosting the response in seropositives. Additionally, we assessed the avidity of the interaction between vaccine-elicited antibodies and the gB immunogen (Fig. 1C) or whole virions (Fig. 1D) as a marker for the strength of the antibody–antigen interaction generated by B cell somatic hypermutation. We identified that the relative avidity index (RAI) of vaccine-elicited antibodies against the gB immunogen was similar to that observed in chronic infection, though somewhat reduced when assessed against whole virions (median RAI: gB/MF59=0.81, SP=0.99; p<0.001, pooled t test).

**Figure 1.**
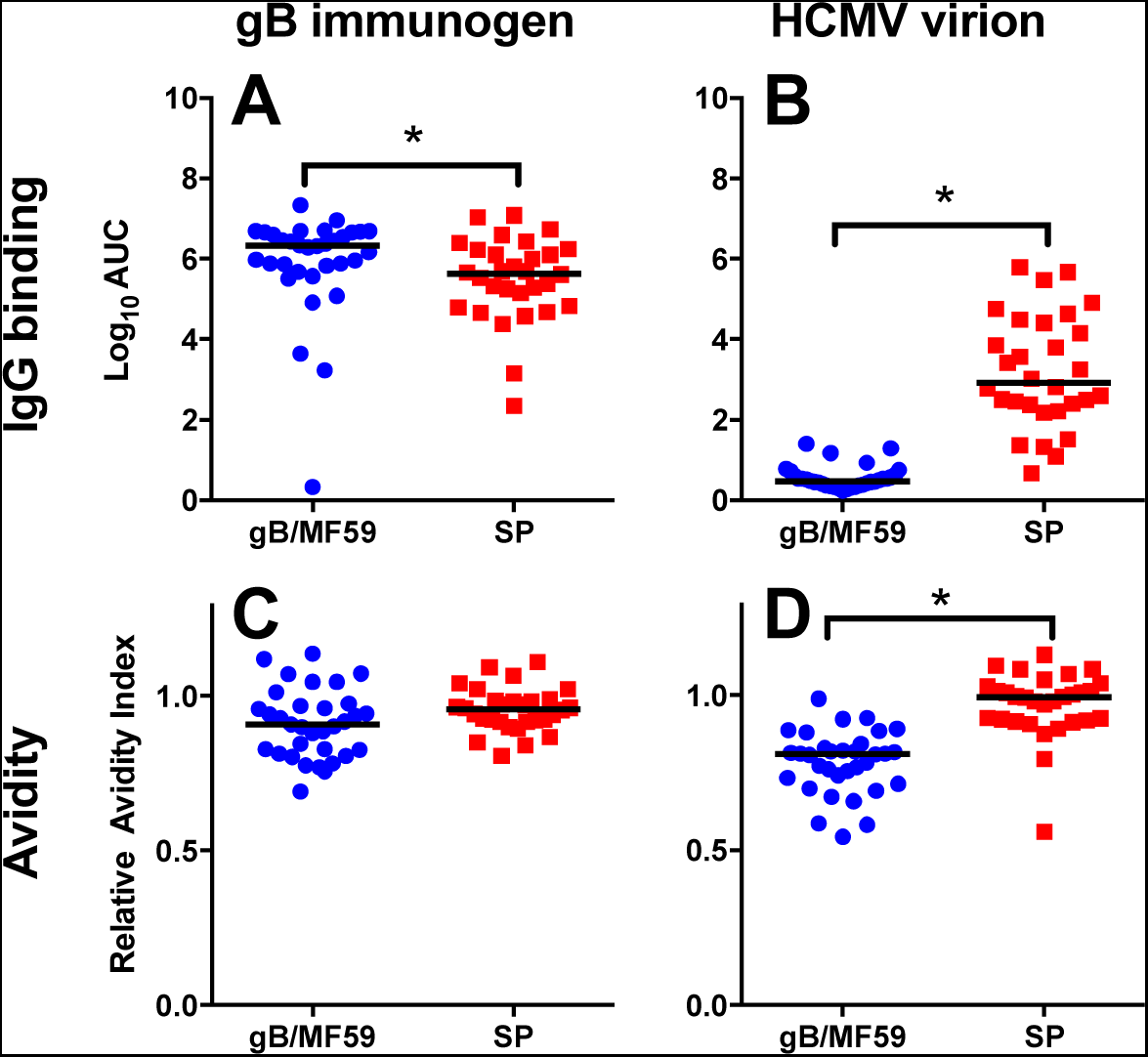
gB/MF59 immunization elicits a robust, high-avidity binding antibody response. The magnitude and avidity of gB immunogen-specific (A,C) and whole HCMV virion-specific (TB40/E strain) (B,D) binding antibodies were assessed for 33 gB/MF59 vaccinees (blue circle) and 30 seropositive, chronically HCMV-infected women (red square). In comparison to the seropositive control cohort, gB/MF59 vaccinees had slightly enhanced gB binding (A) though reduced HCMV virion binding (B). Additionally, in comparison to seropositive women, gB/MF59-elicited antibodies had comparable gB-binding avidity (C), though reduced HCMV virion binding avidity (D). Each data point represents the mean value of 2 experimental replicates. Horizontal values indicate the median values for each group. *=p<0.05, Satterthwaite t test (IgG binding), pooled t test (avidity).

### HCMV neutralization and IgG binding to gB-neutralizing domains

We next investigated the ability of vaccine-elicited antibodies to neutralize a panel of HCMV strains, including the autologous Towne strain (Fig. 2A), AD169 (Fig. 2B), and TB40/E (in fibroblasts, Fig. 2C, and epithelial cells, Fig. 2D). Assays were conducted in both the presence and absence of rabbit complement to assess the possibility of enhanced neutralization titers following complement fixation. A low level of vaccine-elicited neutralization was observed against the autologous Towne virus, though significantly reduced compared to the seropositive group (Towne median log_10_ID_50_: gB/MF59=1.70, SP=2.96; p<0.001, pooled t test). Very few vaccinee samples had detectable neutralization against heterologous viruses, though neutralization of these strains was robust in the seropositive group (TB40/E epithelial cell median log_10_ID_50_: gB/MF59<1; SP=3.80; p<0.001, Fisher’s exact test). We observed minimal enhancement of vaccine-mediated neutralization activity in the presence of complement (Towne median vaccinee log_10_ID_50_: comp=1.70; no comp=1.93), not quite as robust as previous reports (41). Subsequently, we examined whether vaccine-elicited antibodies bound to previously-identified gB neutralizing epitopes (23, 28), including AD-1 (Fig. 2F), AD-2 site 1 (Fig. 2G), Domain I (AD-5) (Fig. 2H), Domain II (AD-4) (Fig. 2I), and Domain I+II combined (Fig. 2J). Intriguingly, though vaccination elicited robust gB-binding responses, there was very poor targeting of these gB neutralizing epitopes. Compared to chronically HCMV-infected, seropositive women, there was significantly reduced vaccine-elicited binding against AD-1 (median log_10_MFI: gB/MF59=1.75, SP=2.46; p=0.001, pooled t test), AD-2 site 1 (median log_10_MFI: gB/MF59=0.65, SP=1.70; p=0.002, Saittertherwaite t test), Domain I (median log_10_MFI: gB/MF59=1.70, SP=3.20; p<0.001, pooled t test), and a Domain I+II fused construct (median log_10_MFI: gB/MF59=2.60, SP=3.14; p<0.001, pooled t test). Thus, the poor targeting of known gB neutralizing epitopes by vaccination likely explains the lack of neutralization responses observed.

**Figure 2.**
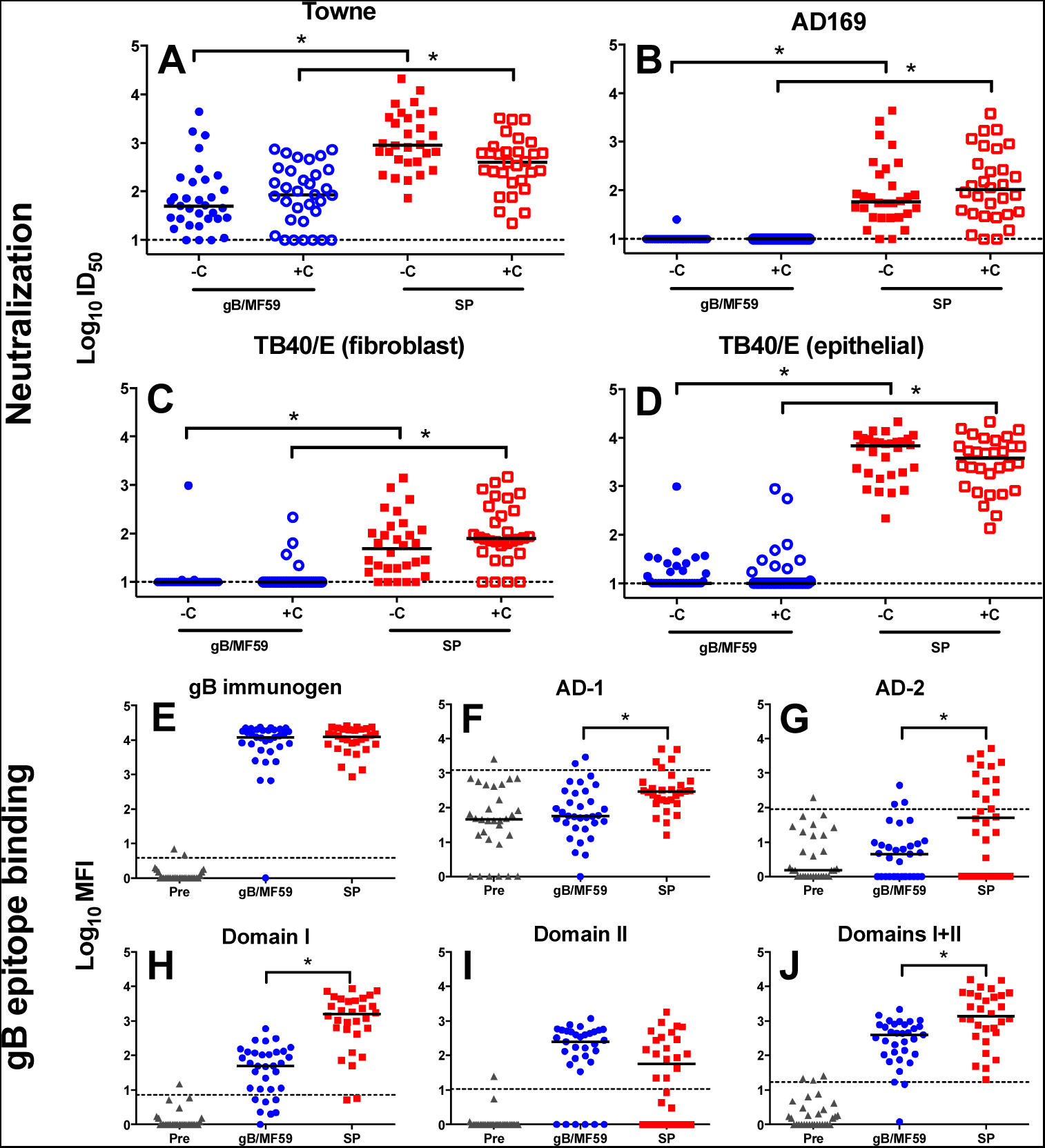
Limited vaccine-elicited neutralization responses and poor gB neutralizing epitope-binding following gB/MF59 immunization. INeutralizing antibody responses (A-D) and gB neutralizing epitope binding (E-J) were assessed for 33 gB/MF59 vaccinees (blue circle) and 30 seropositive, chronically-HCMV infected individuals (red square). Neutralization was measured against Towne strain HCMV (A) and AD169 strain HCMV (B) in fibroblasts, and TB40/E strain HCMV in both fibroblasts (C) and epithelial cells (D). Assays were conducted in both the presence (“+C”, solid symbols) and absence (“-C”, open symbols) of purified rabbit complement. In comparison to seropositive, chronically-infected individuals, gB/MF59 vaccine-elicited neutralization titers were reduced against each viral strain (A-D). Binding responses against the gB immunogen (E) and known gB neutralizing epitopes AD-1 (F), AD-2 (G), Domain I (H), Domain II (I), and Domain I+II combined (J) were measured. In comparison to seropositive women, gB/MF59 vaccination elicited reduced binding against AD-1, AD-2, Domain I, and Domain I+II combined. Each data point represents the mean of 2 experimental replicates. Horizontal dotted lines for neutralization assays indicates the starting dilution, whereas dotted lines for neutralizing-epitope binding indicate the threshold for positivity (preimmune control mean + 2 standard deviations). Black horizontal bars indicate the median values for each group. *=p<0.05, Fisher’s exact test (neutralization), pooled t test (epitope binding).

### Linear gB epitope binding

To map the epitopes targeted by vaccine-induced antibodies a peptide microarray library was created, consisting of 15-mers overlapping each subsequent peptide by 10 residues and spanning the entire gB open reading frame (Towne strain). We observed that vaccine-elicited linear epitope binding was quite distinct from that observed in the setting of chronic HCMV infection (Fig. 3, Fig. S2). Most notably, there was a negligible AD-2 site 1 response elicited by vaccination, which is known to be a target of potent gB-specific neutralizing antibodies (42) (AD-2 site 1 median log_10_MFI: gB/MF59=2.68, SP=3.69; p<0.001, Saittertherwaite t test). Furthermore, vaccination resulted in dominant IgG response against the non-neutralizing AD-3 epitope located in the gB protein cytodomain (AD-3 median log_10_MFI: gB/MF59=5.23, SP=4.14; p<0.001, pooled t test), comprising 76% of the linear gB IgG response in vaccinees compared to 32% in chronically HCMV-infected individuals (Fig. S2A,B,F)

**Figure 3.**
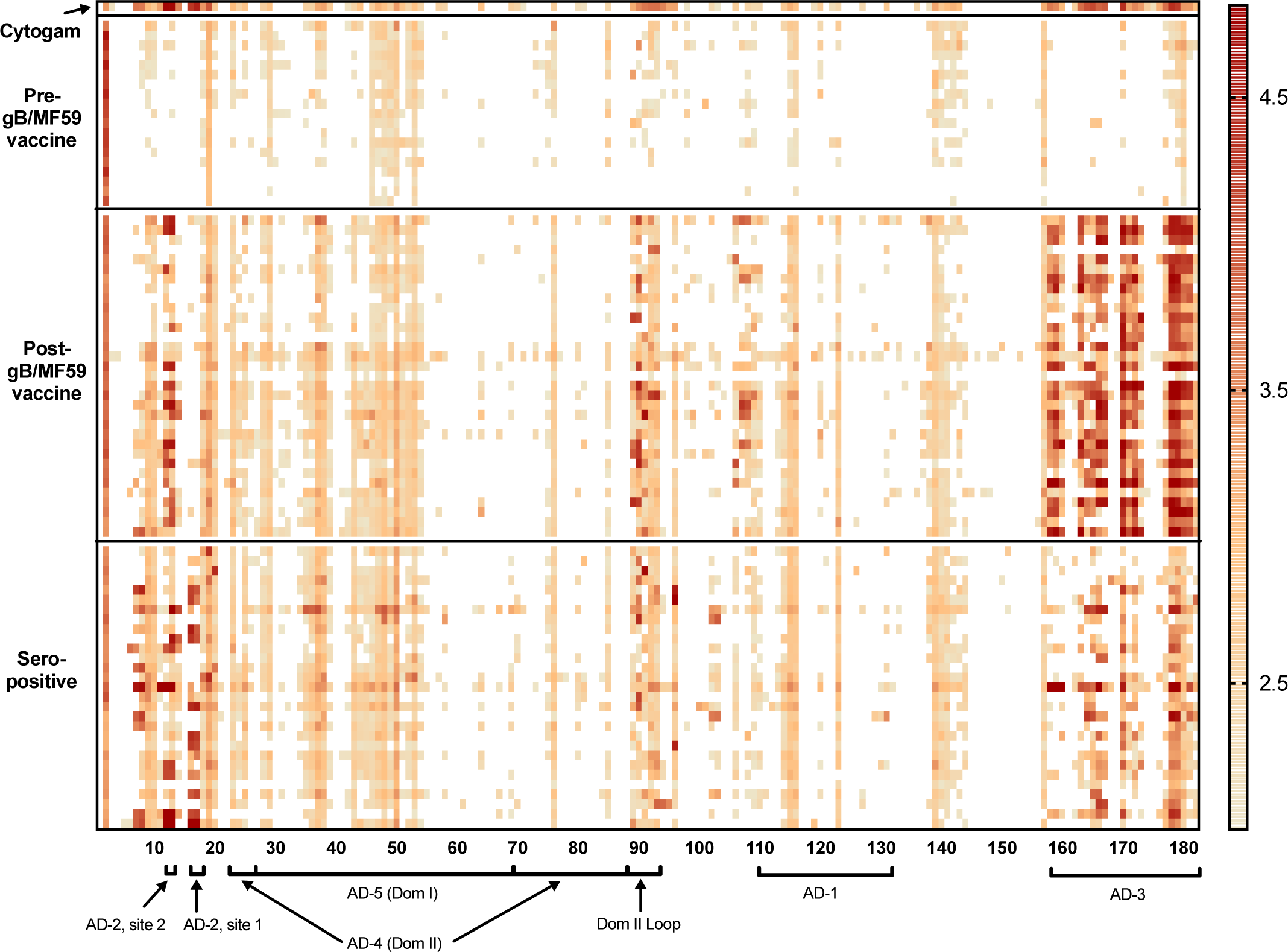
Dominant linear epitope binding response against cytosolic antigenic domain 3 following gB/MF59 immunization. The binding magnitude of antibody responses of Cytogam, 19 gB/MF59 vaccinees pre-immunization, 32 gB/MF59 vaccinees post-immunization, and 30 chronically-infected seropositive controls were assessed against a 15-mer peptide library spanning the entire Towne gB open reading frame (180 unique peptides). Each row indicates a single patient. Assay was completed in triplicate, and the binding magnitude is indicated as the log-scaled, median fluorescent intensity. White indicates median fluorescent intensity < 100. Peptides corresponding to distinct gB antigenic domains are indicated along X-axis.

### IgG subclass distribution and binding to membrane-associated gB

Given the poor neutralizing antibody responses observed, we sought to investigate whether vaccine-elicited IgG responses had properties suggestive of the ability to mediate non-neutralizing antibody effector functions. First, we examined the IgG subclass of gB-directed responses, and identified that both vaccinees and chronically HCMV-infected individuals had a similar response profile dominated by IgG1 and IgG3, with very little detectable IgG2 or IgG4 subclasses (Fig. 4A-D). Furthermore, we examined whether vaccine-elicited immune responses could bind to membrane-associated gB expressed on the surface of both gB-transfected (Fig. 4E,F) and TB40/E-infected cells (Fig. 4 H,I). Vaccine-elicited IgG bound to both transfected autologous Towne strain gB (Fig. 4E) and a heterologous strain gB (Fig. 4F) (most frequently-detected strain in infected vaccinees) more robustly than antibodies elicited by chronic HCMV infection (heterologous median % IgG binding: gB/MF59=13.2%, SP=5.9%; p<0.001, Saittertherwaite t test). Yet, vaccine-elicited IgG bound TB40/E-infected cells less well than antibodies elicited by chronic infection (Fig. 4H) (median % infected cell binding: gB/MF59=6.7%, SP=36.7%; p<0.001, Saittertherwaite t test), likely due to antibody binding to other glycoprotein epitopes in the seropositive group. Thus, we purified gB-specific IgG from both gB/MF59 vaccine and SP plasma and assessed the magnitude of the infected cell-associated gB binding. The purified gB-specific IgG revealed higher magnitude infected cell-associated gB binding in the vaccinee group (Fig. 4I) (median % infected cell gB binding: gB/MF59=4.4%, SP=1.7%; p=0.01, Saittertherwaite T test), consistent with the gB-transfected cell IgG binding magnitude. Finally, we examined NK cell degranulation in the presence of plasma antibodies from gB vaccinees as this process is a prerequisite of both antibody-dependent cellular cytotoxicity (ADCC) and cytokine release by activated NK cells. Interestingly, despite the vaccine eliciting robust gB transfected and infected cell IgG binding, minimal NK cell degranulation responses were detected in vaccinees using both gB-transfected (Fig. 4G) and TB40/E-infected target cells (Fig. 4J). In contrast, the majority of chronically HCMV-infected individuals had antibodies that mediated measurable, though low magnitude, NK degranulation (TB40/E-infected targets, % CD107a+ NK cells: gB/MF59=4.9%, SP=6.6%; p<0.001, Wilcoxon rank sum test).

**Figure 4.**
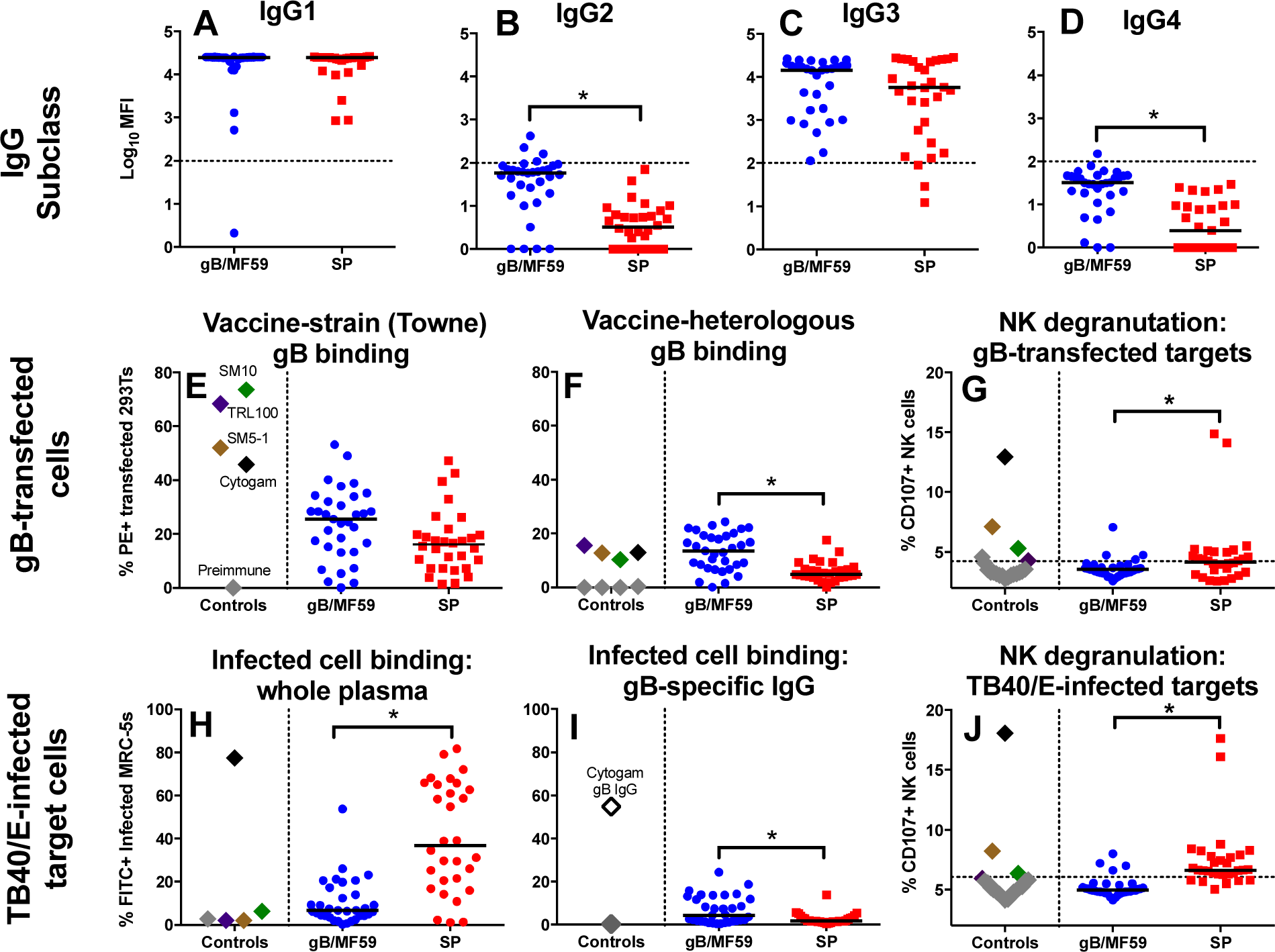
gB/MF59 vaccination elicited high-magnitude IgG3 responses and robust membrane-associated gB IgG binding. The magnitude of gB-specific IgG1 (A), IgG2 (B), IgG3 (C), and IgG4 (D) subclass responses was assessed for 33 gB/MF59 vaccinees (blue circles) and 30 seropositive, chronically-HCMV infected individuals (red squares). A similar IgG1 and IgG3 subclass profile was elicited by both gB vaccination and chronic infection, with nearly undetectable levels of IgG2 and IgG4 gB-specific antibodies. Furthermore, the ability of plasma antibodies to bind to membrane-associated gB expressed on the surface of transfected cells was assessed, including autologous (Towne) (E) and heterologous gB (most frequently identified strain in infected vaccinees) (F). Likewise, binding to TB40/E-infected cells was quantified using both whole plasma (H) and purified gB-specific IgG (I). Lastly, the ability of plasma antibodies to activate NK cells in the presence of either gB mRNA-transfected ARPE target cells (G) or TB40/E-infected ARPE target cells (J) was assessed by the percentage of NK cells expressing CD107a. Black horizontal bars indicate the median values for each group. Horizontal dotted lines in subclass plots (A-D) indicate the threshold for positivity, defined here as 100 MFI. The dotted line in the NK cell degranulation plots (G,J) represents the threshold for positivity (mean of preimmune samples + 2 standard deviations). Nonspecific binding to transfected and infected cells was adjusted for by subtraction of % positive cells against negative control cell population, while nonspecific NK cell degranulation was not corrected. Each data point represents the mean value of two experimental replicates. For E-J, control values are displayed to indicate a dynamic range of the assay, including cytogam (black), AD-2 mAb TRL345 (purple), Dom I mAb SM10 (green), and Dom II mAb SM5-1 (brown). *=p<0.05, Satterthwaite t test.

### Antibody-mediated phagocytosis and monocyte infection

We next investigated the ability of vaccine-elicited antibodies to mediate monocyte phagocytosis. We developed highly-specific, flow-based assays for measuring the phagocytosis of both gB-conjugated beads (Fig. 5A,B) and fluorescently-conjugated HCMV virions (Fig. 5C,D). Cellular uptake of HCMV was confirmed by confocal microscopy, demonstrating fluorescent virus either at the cell surface (Fig. 5E) or internalized (Fig. 5F). The small number of distinct viral foci observed following phagocytosis differs from the multiple, dispersed foci following monocyte infection (Fig. 5H). Additionally, we confirmed that phagocytosis did not lead to productive infection, as cells that phagocytosed TB40/E-mCherry virus did not exhibit mCherry expression at 48 hours post phagocytosis (Fig. S3). Robust vaccine-elicited phagocytosis of gB immunogen-coupled beads was observed, exceeding that in SP individuals (median % phagocytosing cells: gB/MF59=52.1, SP=29.1; p=0.04, pooled T test) (Fig. 5I). To determine whether the dominant AD-3 antibody response (Fig. 3) contributed to this vaccine-elicited gB phagocytosis observed, we investigated phagocytosis of gB ectodomain-coupled beads (Fig. 5J). There was a reduced magnitude of phagocytosis activity directed against the gB ectodomain in comparison to the full gB protein. Furthermore, there was no observable difference in gB ectodomain-directed phagocytosis activity between vaccinees and SP individuals (median % phagocytosing cells: gB/MF59=24.8, SP=18.6; p=ns, pooled t test), suggesting that a proportion of the phagocytosis-mediating antibodies measured in vaccinees target cytodomain epitopes likely not exposed on the surface of an HCMV virion or infected cell. Additionally, we examined phagocytosis of whole HCMV virions and noted more robust virion phagocytosis mediated by plasma antibodies from chronically HCMV-infected individuals compared to vaccinees (Fig. 5K) (median % phagocytosing cells: gB/MF59=9.6%, SP=17.6%; p<0.001, pooled t test). To determine if antibodies targeting other glycoprotein epitopes in the SP group contribute to this difference, we evaluated virion phagocytosis mediated by purified gB-specific IgG from vaccinees and HCMV-infected individuals and observed similar levels of gB-specific phagocytosis (Fig. 5L) (median % positive cells: V=11.3%, SP=10.1%; p=ns, pooled t test). Finally, we confirmed that, though gB vaccine-elicited antibodies could mediate robust virion phagocytosis, they minimally blocked infection of THP-1 monocytes (Fig. 5M,N) (normalized % neutralization at 1:100 dilution: gB/MF59=18.4%, SP=82.8%; p<0.001, Satterthwaite t test).

**Figure 5.**
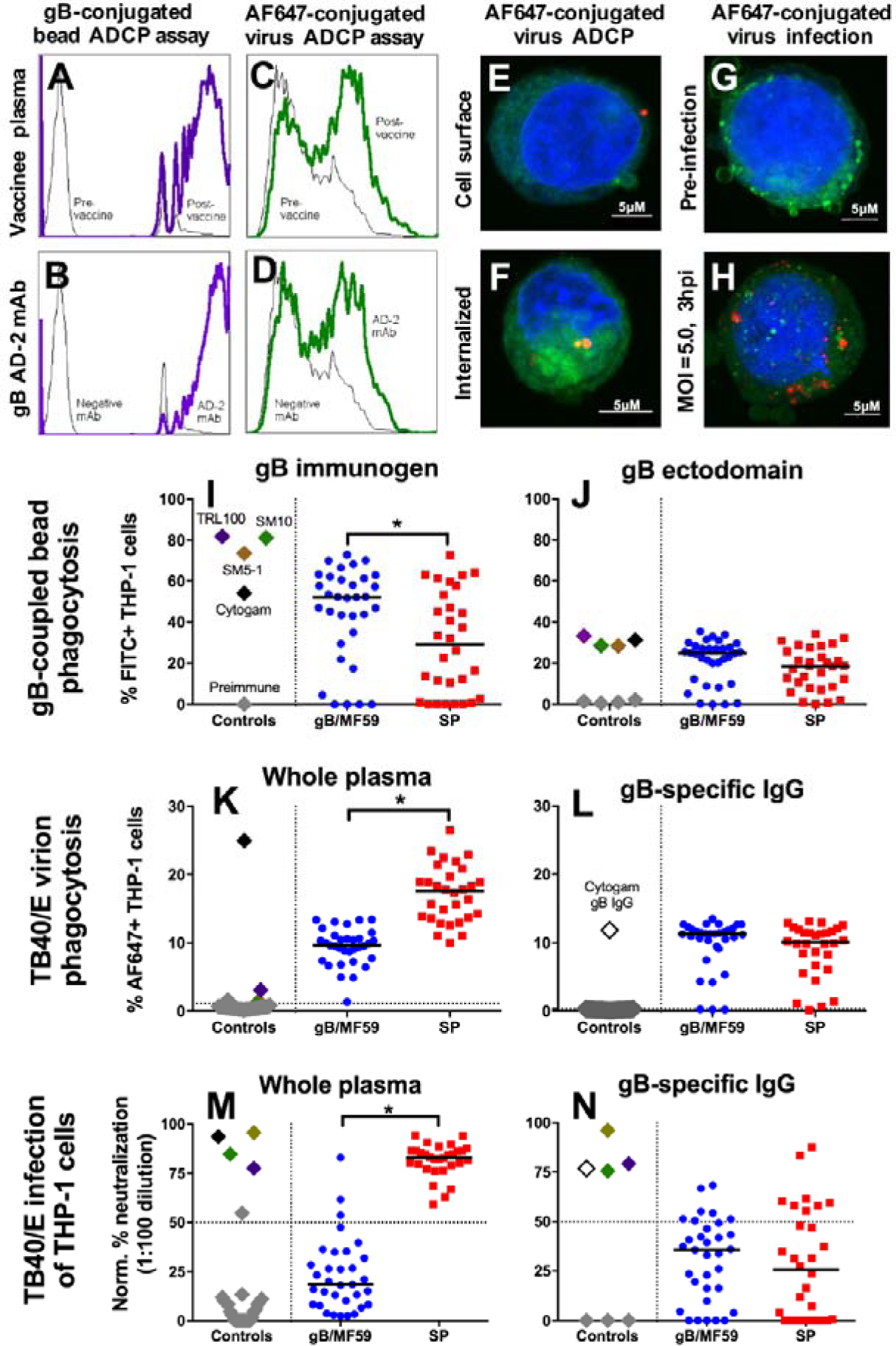
gB vaccine elicits antibodies that mediate robust HCMV virion phagocytosis, though do not block monocyte infection. A flow cytometry-based assay was created to assess antibody-mediated phagocytosis of both gB-coupled fluorescent beads (A,B) and fluorophore-conjugated virus (C,D). Histogram plots of fluorescent intensity indicate the sensitivity of the assay for sera (A,C) and a gB AD-2 specific mAb (B,D). The assay was validated by confocal microscopy (E-G) of THP-1 cells that have either phagocytosed (E,F) or been infected with (G,H) fluorescently-labeled virus. Nuclear material is shown in blue, plasma membrane in green, and AF647-tagged virus in red. Images indicate that phagocytosing cells can either have virus bound to the cell surface (E) or internalized (F). These assays were used to test the phagocytosis-mediating ability of plasma IgG from 33 gB/MF59 vaccinees (blue circles) and 30 chronically HCMV-infected individuals (red squares) of gB immunogen-coupled fluorescent beads (I), gB ectodomain-coupled fluorescent beads (J), and fluorophore-conjugated whole HCMV virions (K,L). In comparison to seropositive, chronically HCMV-infected women, more robust phagocytosis of the gB immunogen and whole HCMV virions (gB-specific activity) was observed among gB/MF59 vaccinees. Lastly, the ability of vaccine-elicited antibodies to block infection of THP-1 cells was assessed at a single dilution (1:100), using both whole plasma (M) and purified gB-specific IgG (N). Black horizontal bars indicate the median values for each group. Nonspecific uptake of fluorescent beads (I,J) was accounted for by subtraction of the % positive cells in the presence of uncoupled beads, while nonspecific uptake of whole virions (C-D) was not corrected. The dotted line in the whole virion phagocytosis plots represents the threshold for positivity (mean of preimmune samples + 2 standard deviations), while the dotted line for the TB40/E infection of THP-1 monocyte plots is the threshold for true neutralization activity (50%). Each data point represents the mean value of two experimental replicates. Control antibody values are displayed to indicate a dynamic range, including cytogam (black), AD-2 mAb TRL345 (purple), Dom I mAb SM10 (green), and Dom II mAb SM5-1 (brown). *=p<0.05, pooled t test.

### Assay correlation matrix

Lastly, we sought to investigate the relationship between measured antibody responses by creating a correlation matrix (Fig. 6). We observed two distinct clusters of responses that appeared related to one another. The first cluster is comprised of gB-specific phagocytosis activity (of both gB protein and whole HCMV virions), binding responses against free gB protein and membrane-associated gB, and gB-specific IgG1 and IgG3 subclass responses. Neutralization activity, by contrast, was associated with a distinct alternate cluster, with robust correlations observed between the different viral strains and cell lines tested. Intriguingly, NK cell activation was inversely correlated with several parameters in cluster 1 (phagocytosis, gB-binding). Finally, some epitope-specific responses correlated with antibody function, including gB domain II-specific IgG responses with phagocytosis activity and AD-2 and domain I-specific IgG responses with neutralization activity. Of note, the formation of cluster 1 was primarily driven by gB-elicited immune responses following vaccination, while cluster 2 was primarily driven by binding and neutralization against non-gB CMV glycoprotein epitopes (Fig. S4A,B).

**Figure 6.**
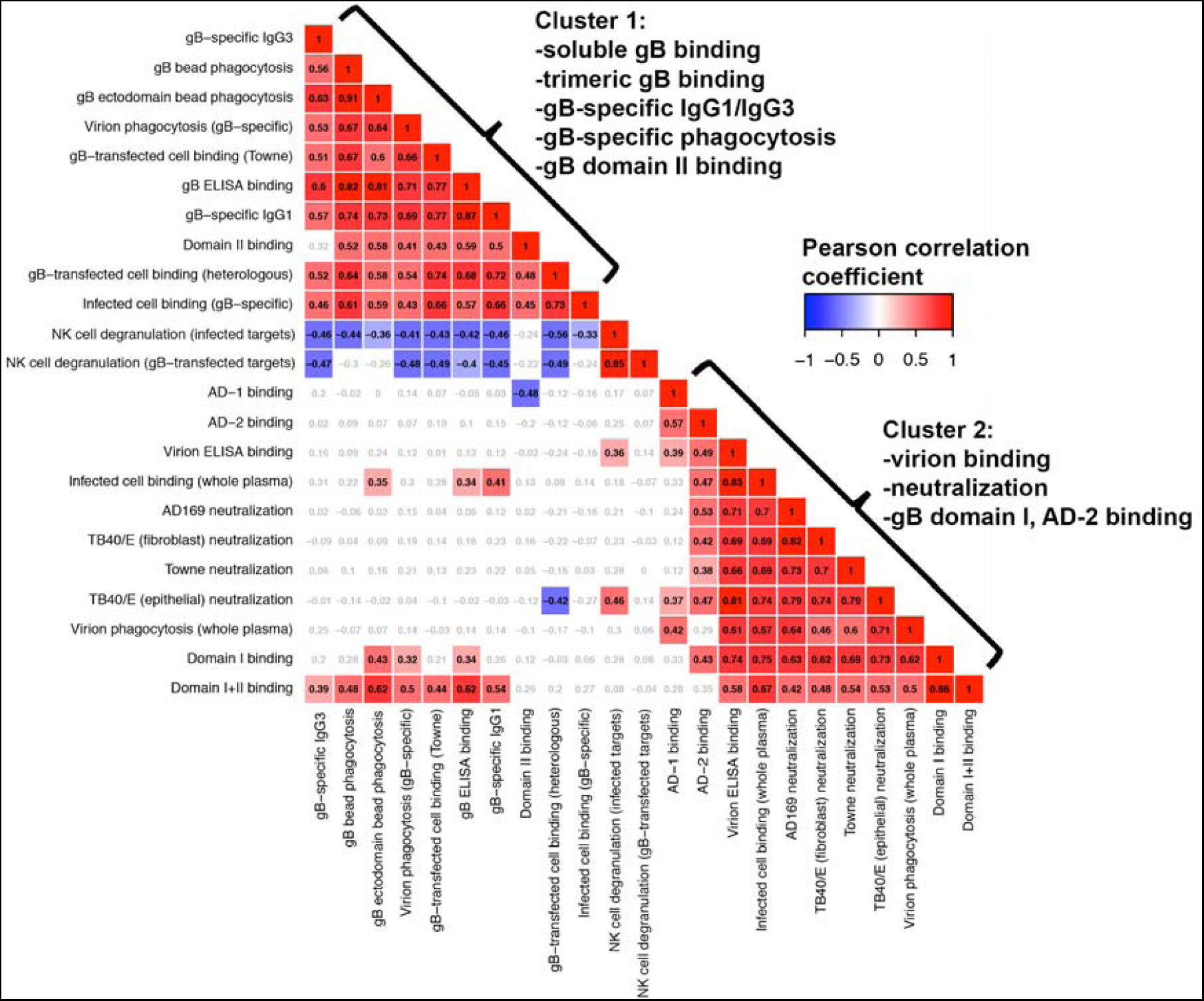
Phagocytosis activity is highly correlated with gB-binding IgG and IgG3 magnitude. A correlation matrix was constructed using data from all 63 tested samples (33 gB/MF59 vaccinee + 30 seropositive) to identify whether assay results correlate with one another. Assays are clustered based on similarity. The Pearson coefficient for each correlation is displayed in the matrix, ranging from −1.0 (blue) to +1.0 (red). Non-significant correlations (p>0.01) are displayed in gray text. Two distinct clusters were identified. The first cluster consists of gB-binding (protein and membrane-associated), IgG1/IgG3 subclass, and ADCP. The second cluster consists of whole virus binding and neutralization activity.

## Discussion

Over the past decade, the HCMV vaccine field has largely shifted its focus away from the elicitation of gB-specific antibody responses and towards the targeting of the gH/gL/UL128/UL130/UL131A pentameric complex (PC) because this protein construct was newly identified as a primary target of potent HCMV neutralizing antibodies (24, 43). Yet, it is important to recognize that the gB/MF59 vaccine platform, which did not include the PC, achieved approximately 50% vaccine efficacy in preventing primary HCMV infection in two phase II trials (14, 15) and demonstrated a protective benefit for transplant recipients (20) without the elicitation of potent neutralizing antibody responses (20, 25, 26). Increasingly, both the limitations of neutralizing antibodies in controlling HCMV cell-to-cell spread (42) and the protective capacity of non-neutralizing HCMV-specific antibodies are becoming recognized (44), indicating that the role of non-neutralizing antibodies needs to be further investigated as a potentially-important endpoint for HCMV vaccine immunogenicity trials.

In this study, we confirmed previous reports of negligible gB/MF59-elicited heterologous and autologous neutralization responses using a panel of viral strains representing diverse gB genotypes: AD169 (gB2), TB40/E (gB4), and Towne (gB1), the autologous virus that is the basis of the vaccine gB antigen (45). While we observed complement-mediated enhancement of HCMV-seropositive neutralization titers in fibroblast cells (2-4 fold), but not in epithelial cells, consistent with previous reports (46). Furthermore, we observed minimal complement-mediated enhancement of gB/MF59 vaccine recipient neutralizing responses, though not as robust as reported previously (41). The limited complement-mediated enhancement is in spite of high levels of gB-specific IgG3 antibodies in the plasma of both vaccinees and seropositive individuals, which is frequently associated with robust complement fixation (47, 48). Congruent with the poor vaccine-elicited neutralizing antibody response, we observed that gB vaccination failed to elicit responses against well-characterized conformational and linear gB neutralizing epitopes such as AD-1, AD-2, Domain I, and Domain II. This finding suggests that these important neutralizing epitopes were either: 1) inadequately exposed to the immune system on the protein immunogen, or 2) that their post-fusion conformation presented via the protein immunogen did not accurately mimic the pre-fusion conformation on the viral envelope, as has been described for HSV-1 gB (49).

Intriguingly, analysis of the linear gB epitope IgG binding profile revealed high-magnitude vaccine-elicited antibody responses against the AD-3 region located within the cytodomain of gB (Fig. 3, Fig. S1) (28, 29). Indeed, an average of 76% of the total vaccine-elicited linear gB IgG binding was directed against this single region, in contrast to 32% in seropositive individuals. Because the AD-3 region is intracellular when gB is expressed on a cell membrane, it presumably does not give rise to antibodies that can bind to or neutralize infectious virus (50). It is unclear whether AD-3-directed antibody responses contributed to vaccine-mediated protection through mechanisms that remain to be defined, or, alternatively, whether this response was merely a diversion away from more functional epitopes. Decoy immune responses away from functional epitopes have been described for other pathogens, most notably for HIV-1, where a vaccine construct containing both gp120 and gp41 elicited a memory B cell response in which 93% targeted the gp41 region (51). This gp41-dominant response is hypothesized to have occurred due to preexisting memory B cells directed against a gp41-crossreactive epitope within intestinal microbiota, thus biasing the antibody response in a form of molecular mimicry. The cause of the immune dominance of the AD-3 region in gB/MF59 vaccinees is unclear, though this epitope may be more accessible on soluble protein compared to its typical intraluminal location on whole virions or HCMV-infected cells. The restricted antibody response against other gB epitopes, including neutralizing epitopes such as AD-1, AD-2, Domain I, and Domain II (Fig. S2) may be a consequence of the high-magnitude linear antibody responses against the AD-3 region. It is therefore possible that a gB vaccine construct without the cytodomain AD-3 epitope would target a greater breadth of epitopes and possibly elicit more potently-neutralizing antibodies.

gB/MF59 vaccination elicited a robust titer of gB-specific IgG3 subclass antibodies, which is unusual for a protein subunit vaccine (52, 53), and possibly due to use of squalene adjuvant MF59 (52). Antigen-specific IgG3 has been implicated in virologic control of other pathogens such as HIV-1 (54), and is anticipated that IgG3 mediates protective antiviral effects by binding to effector cell Fc receptors and facilitating non-neutralizing functions such as antibody-dependent cellular cytotoxicity (ADCC) (55) or antibody-dependent cellular phagocytosis (ADCP) (56). Non-neutralizing antibody effector functions have not been evaluated extensively for HCMV, though NK cells have been strongly implicated in control of HCMV replication (57-60). Moreover, there is some precedent for ADCC-mediated control of HCMV replication *in vitro* in patients with severe AIDS and concomitant HCMV retinitis (61). Furthermore, purified HCMV hyperimmune gamma globulin (Cytogam) has been observed to dramatically enhance the antiviral function of macrophages or NK cells in culture (61). A critical prerequisite for any vaccine-elicited non-neutralizing effector functions is antibody binding to membrane-associated glycoproteins. Importantly, our results demonstrate robust, strain-independent binding of vaccine-elicited antibodies to gB expressed on the surface of both gB-transfected and HCMV-infected cells, suggesting the possibility that these antibodies facilitate antiviral functions such as ADCC or ADCP.

Despite the high-magnitude vaccine-elicited IgG3 response and membrane-associated gB-binding, no substantial evidence of ADCC was observed among vaccinees. Yet, we demonstrated that vaccine-elicited plasma antibodies could mediate a robust level of ADCP of both the gB immunogen alone and gB expressed on the surface of whole virions. Antibody-mediated uptake of whole virions has not to our knowledge been explored as a protective immune mechanism for HCMV, though it has been shown to play an important role in clearing other viral pathogens including influenza (62), West Nile virus (63), adenovirus (64), SARS coronavirus (65), foot-and-mouth disease virus (66), and perhaps HIV-1 (56). As we observed robust vaccine-elicited IgG3 antibody responses, it should be noted that IgG3 has high affinity for the Fc receptors expressed on monocytes/macrophages (48), and that this IgG subclass has been associated with more robust uptake of opsonized virus (56, 67). Monocytes are widely recognized as an important target for HCMV latent infection and dissemination throughout the body (68). Our data suggests that phagocytosed virions are destroyed and do not initiate HCMV replication, yet the fate of a phagocytosed virion is relatively unexplored and subsequent studies should investigate whether antibody-mediated uptake of HCMV can facilitate initiation of latent/lytic HCMV infection.

One limitation of this study is that we did not assess the role of CD4+ or CD8+ T cells in gB/MF59 vaccine-elicited functional immunity, as stored mononuclear cells are not available. Yet protein vaccines, and MF59-adjuvanted protein vaccines in particular, are generally poor at stimulating antigen-specific T cells (69), and previous studies have indicated that MF59 induces a Th2 polarized immune response (70). Nevertheless, this topic remains controversial, as one group has reported robust CD4+/CD8+ T cell immunity in a human cohort following 3 doses of an MF59-adjuvanted protein vaccine (71). Another shortcoming is that we did not have access to a sufficient number of vaccinee samples to compare humoral immune responses between vaccinees who acquired HCMV during the course of the vaccine trial and those who did not. Thus, we cannot say with certainty that non-neutralizing antibody functions such as ADCP were associated with protection against HCMV infection. Nevertheless, this investigation has expanded the repertoire of HCMV-specific antibody functional assays and informs which vaccine-elicited antibody functions were potentially-responsible for the partial vaccine efficacy, guiding subsequent investigation in HCMV vaccine trials.

Some researchers have argued that conventional immune metrics currently used as a surrogate marker of effective anti-HCMV immunity, such as neutralization, may not tell the whole story (72). This in-depth investigation of the immune responses elicited by the most efficacious HCMV vaccine candidate to-date has revealed important biology of antibody functions that are potentially-protective against HCMV infection. Though neutralizing antibody responses are likely important in the control of cell-free HCMV dissemination, cellular responses and/or non-neutralizing antibody effector functions may be essential to eliminate the infected cell reservoir and contain direct cell-to-cell spread. Our data suggest for the first time that gB/MF59-elicited antibodies can mediate robust, non-neutralizing antiviral functions, and that such responses in the absence of potent neutralizing antibodies are the likely mechanism behind the clinically-demonstrated 50% vaccine efficacy. Thus, the elicitation of non-neutralizing antibody responses against HCMV should be a consideration in the rational design of the next generation of HCMV vaccines for the elimination of congenital and transplant-associated HCMV infections.

## Methods

Full methods are available in the supplemental material.

## Author contributions

C.S.N. and S.R.P. designed research; C.S.N., T.H., and E.C. performed research; C.S.N., T.H., E.C., G.X., and N.V. analyzed data; R.F.P contributed samples and expertise; and C.S.N., J.P., and S.R.P. wrote the paper.

## Acknowledgements

The authors would like to recognize Sam McMillan and Shaunna Shen for their assistance with peptide microarray design, data collection, and analysis, as well as Matthew Tay, Derrick Goodman, and Georgia Tomaras for reagents and technical assistance with the phagocytosis assays. Additionally, we’d like to thank Sanofi Pasteur, Merck Vaccines and Trellis Biosciences for the generous gift of research materials. This work was supported by: NIH/NICHD Director’s New Innovator grant to S.R.P (DP2HD075699) and fellowship grant to C.S.N (F30HD089577). The funders had no role in study design, data collection and interpretation, decision to publish, or the preparation of this manuscript. The content is solely the responsibility of the authors and does not necessarily represent the official views of the National Institutes of Health.

